# A scalable, high-throughput neural development platform identifies shared impact of ASD genes on cell fate and differentiation

**DOI:** 10.1101/2024.09.25.614184

**Authors:** Xuran Wang, Matthew Lalli, Urvashi Thopte, Joseph D. Buxbaum

## Abstract

**Background:** Deleterious mutations in hundreds of genes confer high risk for neurodevelopmental disorders (NDDs), posing significant challenges for therapeutic development. Identifying convergent pathways shared across NDD genes could reveal high-impact therapeutic targets.

**Methods:** To identity convergent pathways in NDD genes, we optimized Perturb-seq, a method combining CRISPR perturbation with single-cell RNA sequencing (scRNA-seq), and applied structural topic modeling (STM) to simultaneously assess impact on cell fate and developmental stage. We then studied a subset of autism spectrum disorder (ASD) genes implicated in regulation of gene expression using these improved molecular and analytical approaches.

**Results:** Results from targeting 60 high-confidence ASD risk genes revealed significant effects on neural development. As expected, ASD risk genes impacted both progenitor fate and/or neuronal differentiation. Using STM, we could identify latent topics jointly capturing cell types, cell fate, and differentiation stages. Repression of ASD risk genes led to changes in topic proportions and effects of four genes (*DEAF1*, *KMT2A*, *MED13L*, and *MYT1L)* were validated in an independent dataset.

**Conclusions:** Our optimized Perturb-seq method, combined with a novel analytical approach, provides a powerful, cost-effective framework for uncovering convergent mechanisms among genes involved in complex neurodevelopmental processes. Application of these methods advanced understanding of the impact of ASD mutations on multiple dimensions of neural development, and provides a framework for a broader examination of the function of NDD risk genes.

## Introduction

Autism spectrum disorder (ASD) affects over 1% of individuals (*1*), imposing a substantial toll on those affected and their families. Ongoing exome sequencing studies have identified over 200 genes harboring mutations with large effect sizes on ASD risk (*2*, *3*), with more to be identified. Similarly, even larger numbers of genes have been identified for intellectual disability and developmental delay (IDD) syndromes (*3*, *4*). Genes for ASD and IDD (collectively neurodevelopmental disorders/NDDs) primarily fall into two functional groups: neuronal communication (NC) and gene expression regulation (GER) (*3*). With such large numbers of genes in ASD and IDD, CRISPR perturbation is being used to systematically disrupt these genes and observe the effects on neural development at a high-throughput scale.

Mutations impacting GER genes are likely to have diverse effects on neural development, including impacting cell fate and/or developmental trajectories. Hence, to translate genetic findings into therapeutics, it is essential to elucidate the impact of gene mutations in the complex context of brain development. The use of induced pluripotent stem cells (iPSCs), CRISPR/Cas9, and single-cell RNA-sequencing (scRNA-seq) provides an unprecedented toolkit to interrogate NDD genes at high-throughput in the context of human neurodevelopment. Perturb-seq, a method coupling CRISPR perturbation scRNA-seq, provides a high-dimensional readout of gene function at medium to high throughput (*5–7*) and can be performed in disease-relevant cells, including modeling neurodevelopment (*8*). We and others (*9–15*) have been applying such methods to study autism risk genes in human iPSC and animal models.

We previously combined CRISPR perturbation and scRNA-seq to elucidate the functions of 14 genes linked to ASD risk (*9*). Building on this, we sought to further increase the scale and disease relevance by perturbing dozens of ASD risk genes in cell types relevant to the disorder. We were particularly interested in understanding the impact of mutations in GER genes, which likely impact multiple aspects of neurodevelopment and neuronal function. However, scaling Perturb-seq to include dozens, and ultimately hundreds, of genes is costly, and interpreting the single readout, i.e., scRNA-seq, is especially challenging in neurodevelopmental models, where changes in cell fate and developmental trajectories are both likely to be major determinants of gene expression. Optimizing Perturb-seq and related approaches to address these barriers is critical, given the large numbers of NDD genes being identified and the focus on neural cell development. Here, we leverage cost-effective methods and deploy novel analytical approaches to address these barriers.

We first validated the Human Neuroscience Panel (10x Genomics) as a cost-efficient tool for targeted Perturb-seq, providing means of focused analysis on neurobiologically-relevant genes. Then, to jointly assess changes in both cell fate and developmental trajectories, we made use of structural topic modeling (STM), an unsupervised machine learning method previously used to organize and understand text documents by inferring latent ‘topics’ from documents (*16*, *17*). Considering scRNA-seq data derived from a single cell as a document, genes as words, and gene expression in a given cell as the word count, we use STM to infer latent topics that relate to cell type/cell fate decisions and developmental trajectory. Each sequenced cell can then be assigned a probability as a specific topic, allowing us to relate CRISPR perturbation to even very subtle changes in topic membership.

We successfully applied targeted Perturb-seq and STM to 60 ASD GER genes from a recent analysis (*3*) to investigate convergent mechanisms of ASD risk genes at molecular and cellular levels. We identified genes that impact cell fate, genes that impact developmental trajectory, and genes that impact both cell fate and developmental trajectory. Additionally, we discovered a set of genes that, when perturbed, have a convergent impact on shifts in topic membership. We observed overlapping findings when we analyzed a published, independent sample that incorporated additional cell types. Our approach provides a means of assessing the impact of mutations in NDD risk genes in a relevant dynamic context, including identifying converging pathways, which can guide the development of targeted therapies and interventions.

## Methods and Materials

### iPSC Culture and NPC Differentiation

#### Cell Line and Maintenance

WTC11 iPSCs with dCas9-KRAB inserted into the second intron of the CYLBL safe harbor locus were obtained from the Allen Institute for Cell Science via Coriell (AICS-0090-391) and maintained according to the AICS standard operating procedure (SOP).

#### Differentiation Protocol

iPSCs were dissociated using Accutase and plated at high density (2 million cells per well of a 6-well plate) on Matrigel or Geltrex in StemFlex or mTesR with ROCK inhibitor (Thiazovivin, 2 µM). The next day, media was changed to include dual inhibitors of SMAD signaling, LDN193189 (100 nM) and SB431542 (10 µM), and the tankyrase inhibitor XAV939 (2 µM) (LSB+X conditions). Over the first 7 days, media was gradually transitioned to neural induction media (NIM) consisting of DMEM/F12 supplemented with 1X N2 and LSB+X. After 7 or 8 days, cells were dissociated into single cells and replated at either 1:1 or 1:2 in NIM with LSB. XAV939 was removed at this point. NPC induction continued for another 7 days without passaging, then cells were replated 1:2 in STEMdiff Neural Progenitor Medium (Stemcell Technologies) for NPC expansion.

### Single-cell RNA-seq Experiment and Analysis

#### Library Preparation

Two biological replicates of independently differentiated NPCs were transduced with separate batches of pooled sgRNA lentivirus. Cells were selected for sgRNA expression by puromycin and grown under self-renewal conditions for two weeks to enable sufficient time for CRISPRi-mediated knockdown. Both replicates of NPCs were prepared in parallel for single-cell RNA-sequencing experiments. Each replicate was loaded across four lanes of a 10x Genomics Chromium Next GEM Chip G at around 10,000 cells per lane. Single-cell libraries were prepared following the Chromium NextGEM single cell v3.1 User Guide (PN-1000121).

#### Sequencing and Processing

Sequencing data corresponding to single-cell transcriptomes were processed using the 10x Genomics software package Cell Ranger. We implemented the Seurat (v 4.0.5) (*18*) workflow for quality control, normalization, scaling, dimensionality reduction, and finding marker genes between clusters. Dimensionality reduction, UMAP visualization, cluster assignment, and pseudotime trajectory were implemented using Monocle3 (v 1.3.7).

### Target Capture

The Human Neuroscience Panel developed by 10x Genomics, targeting 1,186 genes relevant to neurobiology, was used according to the manufacturer’s instructions. The panel was augmented with the ASD risk genes not already included, as well as sequences corresponding to Cas9 and the lentiviral vector.

### sgRNA enrichment

An enrichment PCR based on the CROP-seq protocol was used to identify sgRNAs in scRNA-seq libraries (*9*, *19*). Initially, two sgRNA-specific enrichment PCR replicates were performed for each single-cell library using 1 µL of the single-cell post-cDNA amplification product as a template. A single-step PCR reaction was used to amplify gRNA from total captured cDNA libraries using custom primers in a 25-µL reaction volume with KAPA HiFi HotStart ReadyMix (2X, Roche).

Additional enrichment libraries were constructed using an optimized hemi-nested 2-step enrichment PCR. A customized Python script was used to map sgRNAs to cell barcodes, extracting and filtering cell barcodes from the sequencing reads. For each sgRNA, reads were counted, requiring at least 20 reads per UMI and 5 UMIs per sgRNA. For analysis, only cells with a single sgRNA were retained.

### Knock-down Effect

Knock-down effects were confirmed using the MIMOSCA pipeline(*5*), which outputs the Beta coefficients for each sgRNA. Z-normalized coefficients were used to show on-target repression of genes of interest.

### Statistical Analysis

Structural Topic Model (*16*) was used to analyze CRISPR RNA-sequencing data from the shared 1050 genes and 20 shared targets and the control cells in both iNPC and organoid data from Li et al. (*11*). The `stm` function from stm(1.3.6) package in R (version 4.2.2) was used with 10 topics for iNPC dataset and 20 topics for organoid data. Parameters included `content = ∼ target + batch` and `prevalence = ∼ target + batch`. Differences in averaged topic proportions for targets and non-targeting samples were clustered to generate the tree structure with `hclust`.

## Results

### Targeting of High-risk ASD Genes

ASD risk genes are expressed in multiple cell types during neural cell development, including neural progenitor cells (NPC) and excitatory and inhibitory neurons (18, 19). In addition, previous studies have highlighted the importance of the forebrain in ASD pathology (*20*, *21*). We implemented a forebrain NPC-directed differentiation protocol in a monolayer format (*22–24)* and confirmed the expression of SOX, NESTIN and DAPI in NPCs with immunofluorescence (**Figure 1A** and **1B**).

**Figure 1.**
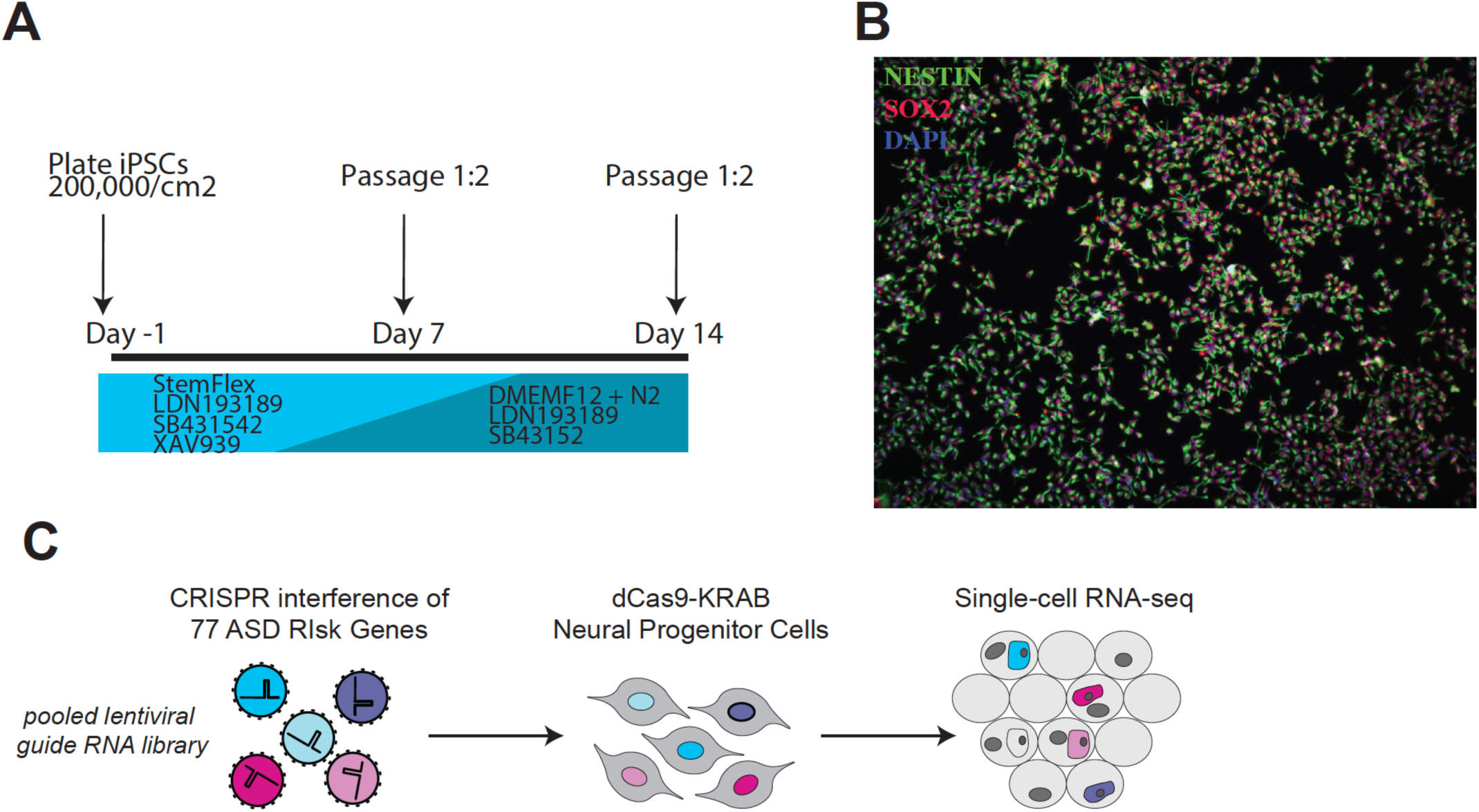
Pipeline of CRISPR interference (CRISPRi) experiment. **A.** Culture timeline of iPSCs differentiation into neural progenitor cells (NPCs). **B.** Immunofluorescence validation of NPC markers. Cells were stained for NESTIN (green), SOX2 (red), and DAPI (blue), confirming the expression of these neural progenitor markers. **C.** Workflow of the CRISPRi experiment targeting 77 high-risk ASD genes. A pooled lentiviral guide RNA (sgRNA) library was used to transduce iPSC-derived NPCs expressing dCas9-KRAB. Following antibiotic selection, single-cell RNA sequencing (scRNA-seq) was performed to assess gene expression changes.

We selected 77 high-confidence ASD risk genes to evaluate their impact on neural differentiation (**Supplementary Table 1**), using CRISPR interference (CRISPRi) (*25*) in an iPSC line containing the CRISPRi machinery integrated into a safe harbor locus (*15*).. In addition to the 58 GER genes described previously (*3*) — chosen due to their mutations having potential impact on both brain development and neuronal function, we included a number of well-studied ASD genes chosen because they represent well-studied genes (e.g., FMR1, MECP2, PTEN, SHANK3, and TSC1), and/or they are amongst the genes commonly impacted by de novo mutations in recent genome-wide studies (e.g., DYRK1A, PTEN, SCN2A, SHANK3, and SYNGAP1), and/or represent additional GER genes implicated in ASD (e.g., AUTS2, CTCF, EHMT1 and KMT2A). sgRNA sequences were designed and cloned individually into a lentiviral vector optimized for sgRNA expression and single-cell capture (*19*, *26*, *27*). To perform perturbation experiments, we transduced iPSC-derived NPCs with a library of sgRNAs targeting 77 ASD risk genes (3 guides each) and 6 non-targeting sgRNAs (237 sgRNAs total) at low multiplicity of infection so that most cells receive 0 or 1 sgRNA. After antibiotic selection of sgRNA-expressing cells, NPCs were grown in self-renewing conditions for 14 days to enable adequate time for CRISPRi repression of target genes to elicit downstream consequences. Single-cell gene expression profiling was performed across two biological replicates of NPCs after ASD risk gene repression using droplet-based scRNA-seq (*28*) (**Figure 1C**; **Supplementary Table 2**). This dataset is referred to as the iNPC (induced NPC) dataset.

### Targeted Single-cell RNA-sequencing

While increasing the number of sgRNAs is straightforward for studying multiple targets simultaneously, Perturb-seq experiments become cost-prohibitive as perturbations increase (*5*). Targeted sequencing of preselected gene panels can potentially address this bottleneck (*29*, *30*). We utilized the Human Neuroscience Panel for target enrichment of single-cell RNA sequencing libraries (10x Genomics) (*28*), augmented with our chosen ASD risk genes. Target capture performed after standard scRNA-seq library preparation enables direct comparison between full transcriptome and target library sequencing. In target capture libraries, 80% of mapped sequencing reads corresponded to the preselected target gene panel, compared to 3% in untargeted libraries. This enrichment approached the theoretical 20-fold improvement predicted by the target capture approach (**Supplementary Figure 1**). At 1/6th of the sequencing depth of the full transcriptome, the target capture library yielded ∼3.5-fold more reads in the target panel (**Table 1**).

**Table 1.** Comparison of sequencing metrics between full transcriptome sequencing and targeted capture sequencing libraries. Targeted capture significantly enriches target reads, providing higher mean target reads per cell and median targeted UMI counts per cell, thus enhancing cost efficiency.

| Metrics | Full Sequencing, Only Targets | Target Capture |
| --- | --- | --- |
| Reads | 667,256,617 | 106,915,094 |
| Mean Reads Per Cell | 63,657 | 10,338 |
| Mean Target Reads Per Cell | 2367 | 8274 |
| Median Target Genes Per Cell | 282 | 364 |
| Total Target Genes | 1204 | 1211 |
| Median Targeted UMI Counts per Cell | 778 | 1,017 |

### CRISPRi Efficiency

Enrichment PCR based on the CROP-seq protocol followed by next-generation sequencing (*9*, *19*) identified sgRNAs in scRNA-seq libraries. Analysis was restricted to cells with only a single identified sgRNA yielded 20,488 (out of 58,958) cells after quality control. Each target gene in our library was covered by a median of 242 cells, and each sgRNA was identified in a median of 83 cells. Reduced expression of targeted ASD risk genes confirmed on-target impact of sgRNA (**Supplemental Figure 2**). Knock-down effects were consistent across replicates and individual sgRNAs, so target-gene level grouping of cells was used throughout analyses. Altogether, 60/77 ASD risk genes were effectively targeted, defined as z-score of the normalized gene expression for non-targeting and perturbations (**Supplementary Figure 2**).

### Perturbation Effects on Neural Cell Development

We elucidated the functional effects of these perturbations on both cell-fate specification and neuronal differentiation. Marker gene analysis (i.e., *SOX2, NES, FOXG1*) confirmed cells as forebrain neural progenitors (**Figure 2; Supplementary Figure 3**). Cells are clustered into 3 main populations (**Figure 2A**). The iNPCs expressed markers of ventral fate (*ASCL1, DLX2*) and GABAergic neuron markers (*GAD1, GAD2*). Two clusters expressed markers of regional NPC identity (i.e., *NKX2-1* and *NKX6-2)* corresponding to intermediate neural progenitor cell NPCs of the medial and caudal ganglionic eminences (MGE-INP and CGE-INP) (**Figure 2B** and **2C**) (*31*). GABAergic interneuron progenitor identity was supported by additional markers (**Supplementary Figure 3**) (*32*).

**Figure 2.**
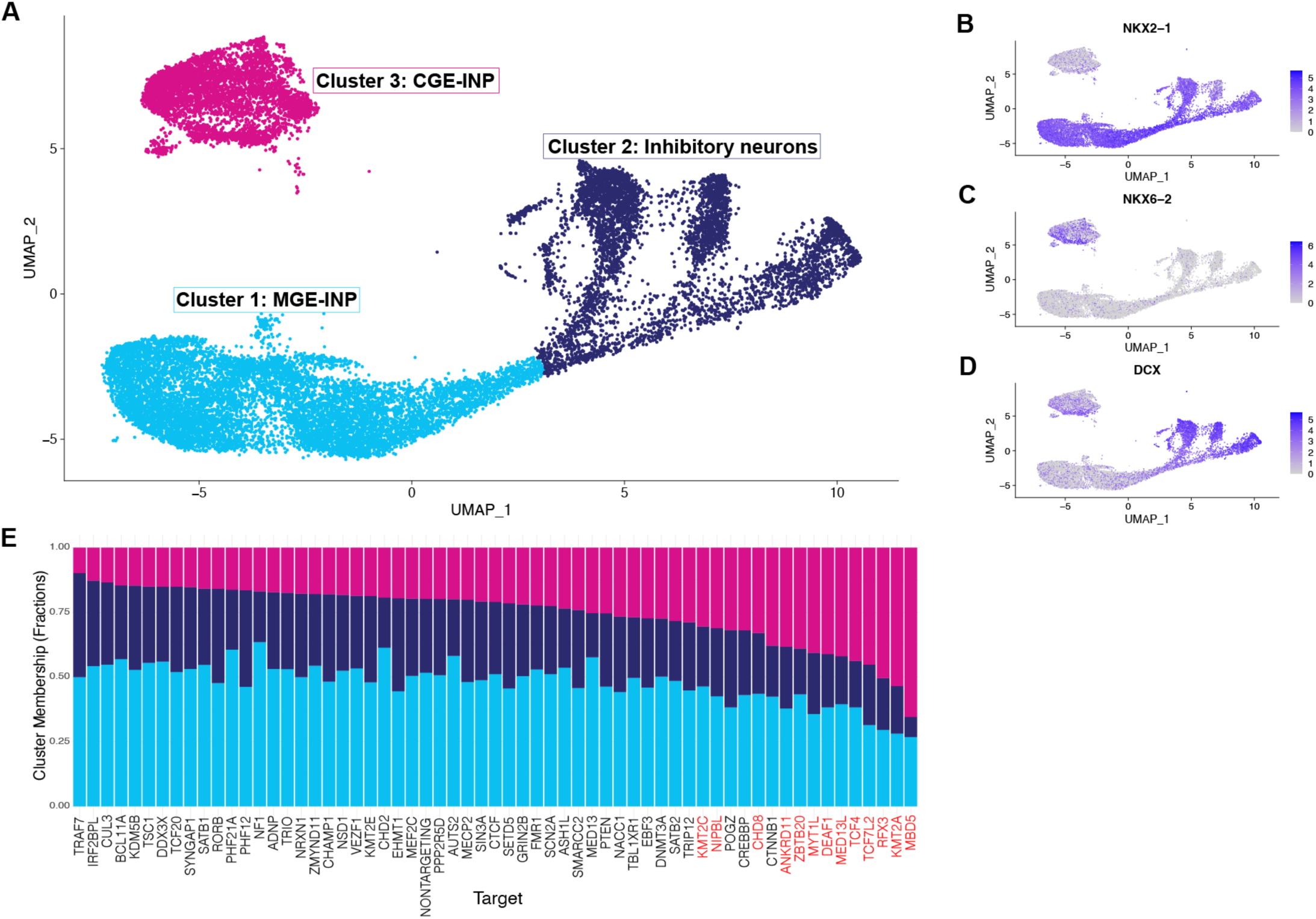
Summary of the iNPC dataset with effective gene expression. **A.** UMAP visualization of the iNPC dataset, with cells colored by clusters. The three identified clusters are Cluster 1 (MGE-INP), Cluster 2 (Inhibitory neurons), and Cluster 3 (CGE-INP). **B-D.** Feature plots showing the expression of regional NPC identity genes. **B.** Expression of NKX2-1; **C.** Expression of NKX6-2; **D.** Expression of DCX. **E.** Barplot showing the fractions of each cluster for the different targets. Targets colored in red significantly differ in cluster fractions compared to the non-targeting cells.

Examining the effects of ASD risk gene perturbation effects on cell-fate specification revealed perturbation impacting cell fate decisions (**Figure 2E**). A set of 13 ASD risk genes, when silenced, significantly increased the proportion of cells identified as CGE-INP (Chi-squared test of CGE-INP vs other cells). These genes include *ANKRD11, CHD8, DEAF1, KMT2A, KMT2C, MBD5, MED13L, MYT1L, NIPBL, RFX3, TCF4, TCF7L2,* and *ZBTB20* (**Supplementary Table 3**).

In parallel, based upon our findings and those of other groups, we hypothesized that some ASD gene mutations alter the pace of neuronal differentiation, with some genes accelerating and others delaying this process (*3*, *9*). Cells from the MGE-INP and inhibitory neuron clusters were ordered along a pseudotime trajectory (**Figure 3A**), corresponding to the progression of neuronal differentiation (**Figure 3B**). The average pseudotime z-score across all sets of cells grouped by their target gene indicate that certain ASD genes appear to impact the pace of neuronal differentiation (**Figure 3C**), either decreasing (e.g., *NF1*, *MED13*, and *MBD5*) or increasing (e.g., *TRAF7* and *POGZ*) the pace (**Figure 3D**). Results using the Kolmogorov-Smirnov test (*33*) for pseudotime density between targeted cells and non-targeting cells also demonstrates changes in pace of differentiation (**Figure 3E**).

**Figure 3.**
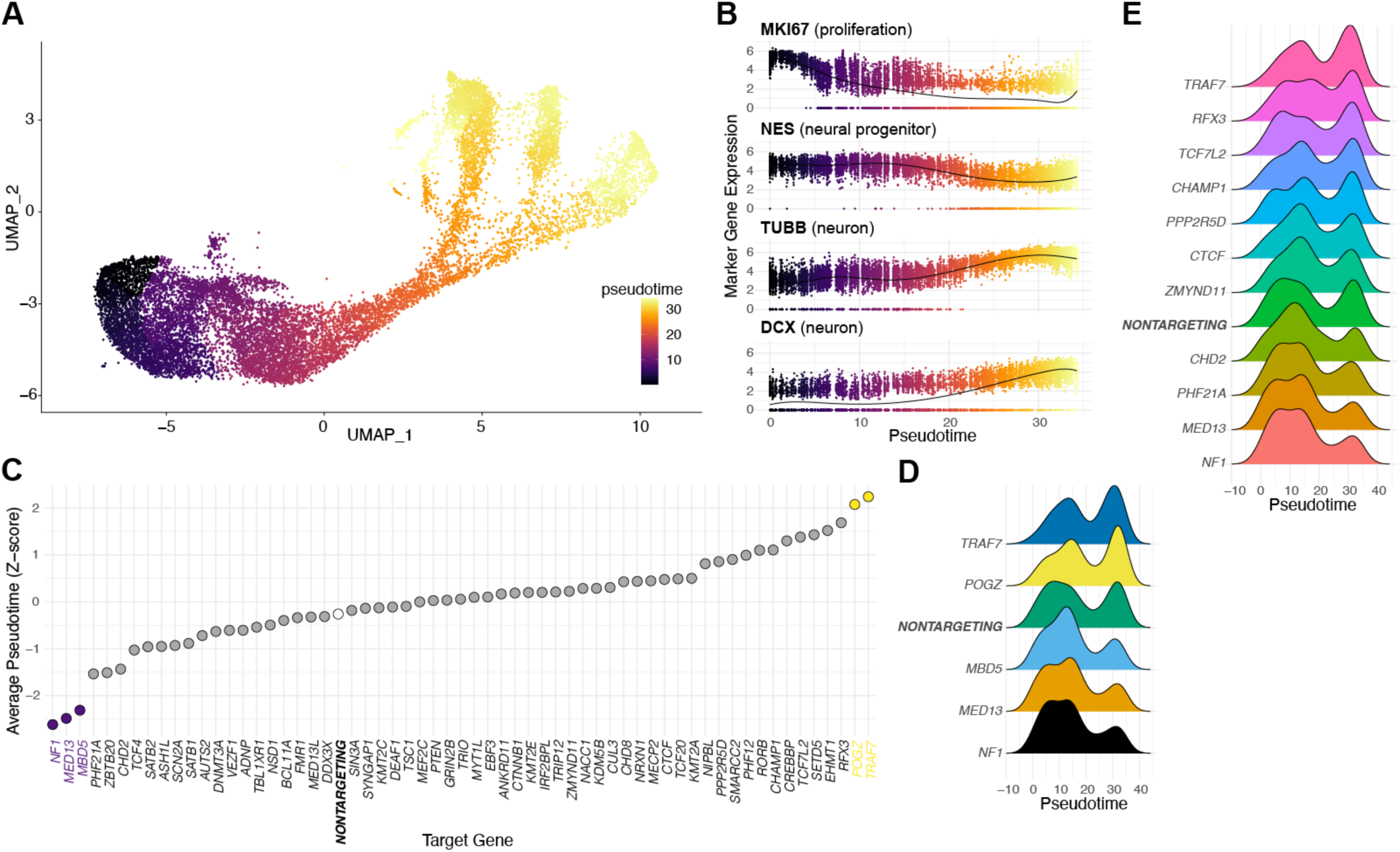
Pseudotime trajectory analysis of the iNPC dataset. **A.** UMAP visualization of Cluster 1 (MGE-INP) and Cluster 2 (Inhibitory neurons), colored by pseudotime as clustered by Monocle3 (*69*, *70*). **B.** Scatter plot showing the expression for marker genes (*MKI67, NES, TUBB*, and *DCX*) across pseudotime, highlighting their roles in proliferation, neural progenitor state, and neuronal differentiation. **C.** Z-score of pseudotime for each target gene, ordered by their pseudotime shift. Significant targets with decreased pseudotime are colored purple, and those with increased pseudotime are colored yellow. **D.** Pseudotime density plots comparing non-targeting cells and cells with significant gene targets (*TRAF7, POGZ, MBED5, MED13*, and *NF1*), showing the distribution of cells along the pseudotime trajectory. **E.** 11 targets shows significant shift based on the Kolmogorov-Smirnov test (K-S test). Among them, TRAF7, NF1, and MED13 also show significant changes in Z-score test. The 11 targets selected by K-S test also show large or small z-scores.

### Structural Topic Modeling

Existing analytical methods for single-cell CRISPR screening often fail to address cell composition heterogeneity, impact on cell fate and differentiation, and the challenges posed by noisy and sparse data (*5*, *34–39*). To capture the full spectrum of cellular response to genetic variations, we applied the structural topic model (STM) to the RNA-seq data, with 10 topics and perturbations as covariates. In our approach, STM assigns weights of genes to each topic, and for each sequenced cell, STM estimates topic proportions. When applying STM to our data, topic proportions are aligned with the cell types and cell clusters identified above (**Supplementary Figure 4A and 4B**). For example, Cluster 3, which is discrete from other clusters (**Figure 2**), is captured by topics 2, 4 and 7. Additionally, topic proportions can capture pseudotime trajectory (**Figure 3**). For example, the more developed neuronal cells (Cluster 2) are captured by topics 1, 6 and 10, while cells earlier in development (Cluster 1) are captured by topic 5, 8, and 9.

Topic proportions across perturbations can be clustered or grouped (**Supplementary Figure 4C**), consistent with convergent effects of perturbing different genes. In these analyses, we observed a significant enrichment of topic 2 with perturbation of *MBD5, KMT2A*, and *RFX3*. These same genes are also implicated in analyses of cell fraction composition (**Figure 2**). Four genes (*NF1, MED13, PHF21A*, and *CHD2*) that show differences in pseudotime analyses (**Figure 3C, 3D,** and **3E**) are members of topic 10.

We also used the difference between average topic proportion between targeted cells and non-targeting cells to capture the perturbation effects on cell membership shift. The convergences across targets are calculated as distance between the cell membership shift vector. For example, the convergences across 60 effective targets in iNPC dataset are shown in **Figure 4A**, indicating shared impacts on cell membership distribution.

**Figure 4.**
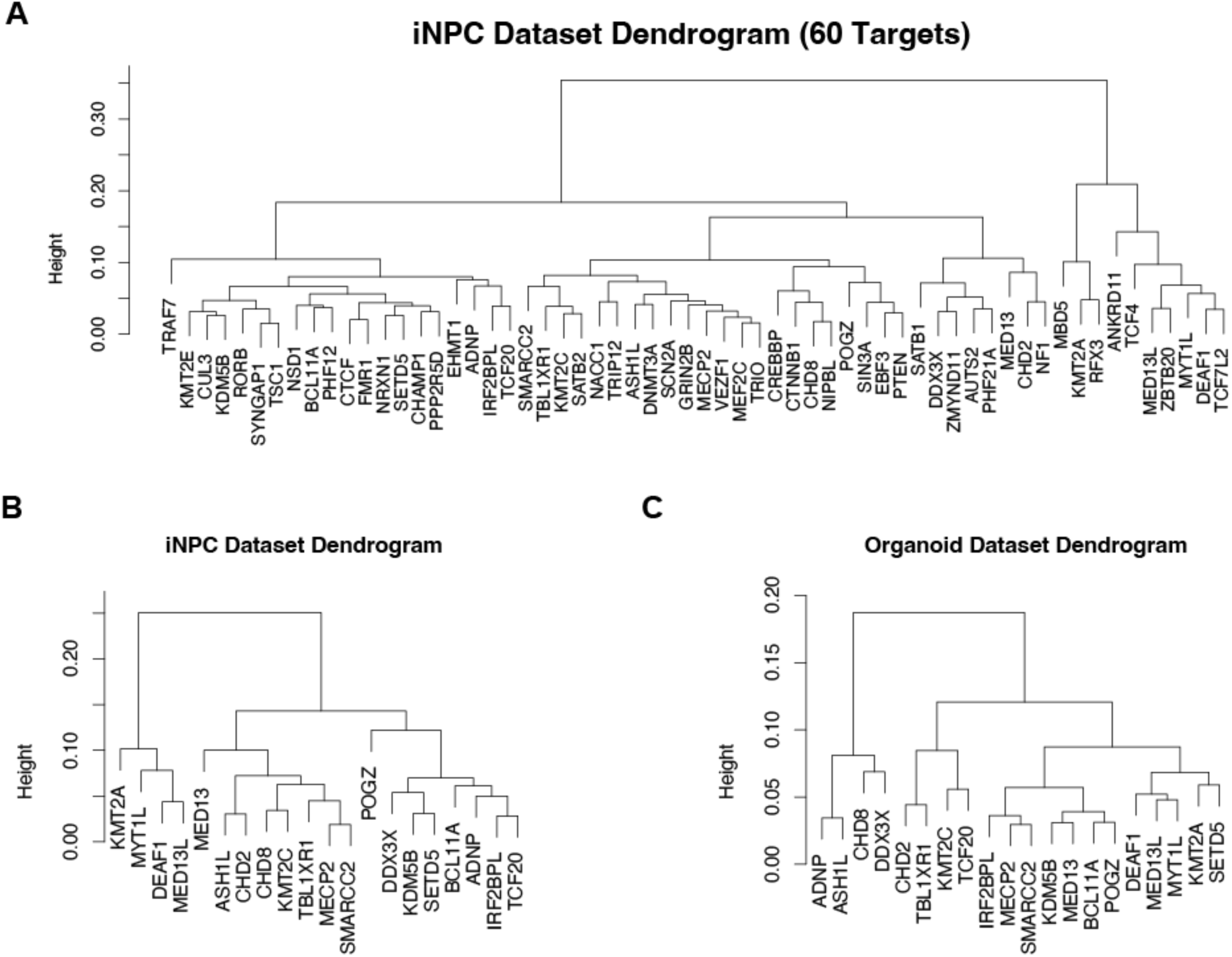
Dendrograms of ASD risk gene targets. These dendrograms illustrate the hierarchical clustering of ASD risk gene targets based on the average difference in topic proportions between control and targeted cells. **A.** Dendrogram of 60 effective targets from the iNPC dataset. **B.** Dendrogram of 20 common targets from the iNPC dataset. **C.** Dendrogram of 20 common targets from the organoid dataset. These dendrograms highlight the clustering patterns of the gene targets, revealing potential similarities and convergent mechanisms in their effects on cellular processes.

### Cell Membership Shift in an Independent Dataset

To validate gene perturbation effects on cell membership and cell fate, an independent dataset from Li et al. (*11*) was examined with STM. This is a single-cell CRISPR RNA-sequencing data of human brain organoids cultured for 4 months, targeting 35 ASD risk genes. The organoid data includes more cell types than in our study, including excitatory neurons, inhibitory neurons and glia cells. (**Supplementary Figure 5A and 5D**). There are cell fraction changes between cell lineage (**Supplementary Figure 5B and 5E**) and between cell states (**Supplementary Figure 5C and 5F**). Focusing on neuronal cells, the pseudotime of cells are also affected by perturbations. Similar to our data, analysis of average pseudotime across all sets of cells indicates that certain ASD genes act by altering the pace of neuronal differentiation (**Supplementary Figure 6**).

As above, we next used the difference between average topic proportion between targeted cells and non-targeting cells to capture the perturbation effects on cell membership shift. We focused on the shared 20 targets from our iNPC data and the published organoid data and first noted that the down-sampling of ASD targets from 60 effective targets to 20 shared targets does not change the tree structure (**Figure 4A** and **4B**), indicating the robustness of the STM in detecting cell membership shifts. Comparing the tree structures from two independent datasets, the ASD risk genes, *KMT2A, MYT1L, DEAF1, and MED13L,* shared similar topic proportion changes (**Figure 4B** and **4C**).

Returning to the iNPC dataset, we further investigate the average topic proportion changes between targeted and non-targeting cells ( **Figure 5A**). We found that the ASD risk genes, *KMT2A, SETD5, DEAF1* and *MED13L*, shared similar topic proportion changes with an increase in Topic 8 and Topic 2 and a decrease in Topic 3, 6, 5, and 9. Using the log fold change to calculate topic specificity, we identified topic-specific genes for the iNPC data (**Supplementary Table 4**). We observed the GO terms for Topic 2 are for the regulation of the development process while Topic 3 GO terms include the negative regulation of the developmental process.

**Figure 5.**
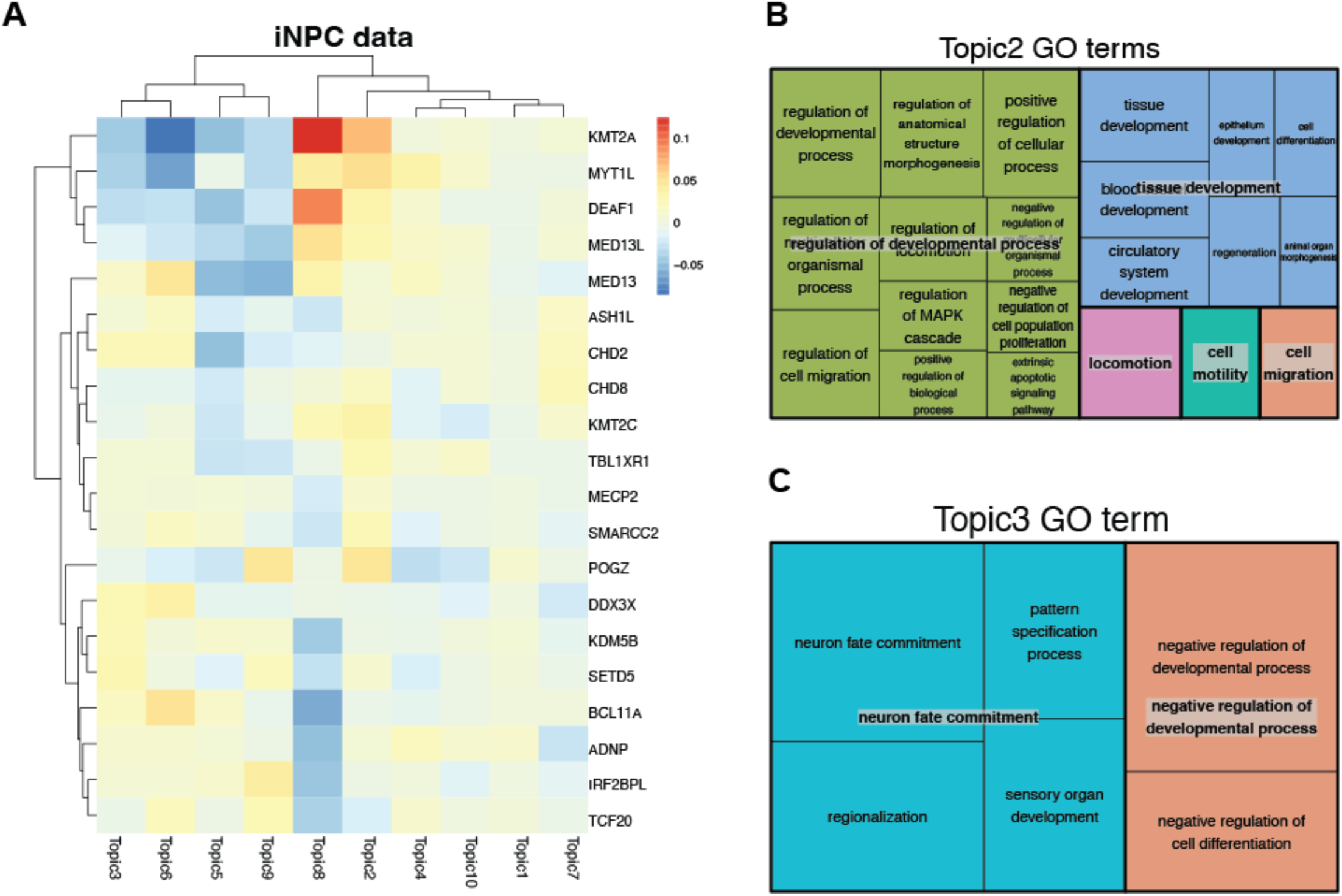
Topic-specific analysis of ASD risk gene targets. **A.** Heatmap showing the shift in topic proportions between targeted and non-targeting cells for various ASD risk genes in the iNPC dataset. The color scale represents the magnitude of the shift, with red indicating an increase and blue indicating a decrease in topic proportion. **B.** Gene Ontology (GO) terms associated with Topic 2 in the iNPC dataset, highlighting biological processes such as regulation of developmental process, tissue development, and cell migration. **C.** Gene Ontology (GO) term associated with Topic 3 in the iNPC dataset, focusing on processes like neuron fate commitment, pattern specification process, and negative regulation of developmental process.

## Discussion

We and others (*3*, *40*) have estimated that approximately 1000 genes carrying rare deleterious mutations contribute to ASD risk. This estimate aligns with the rate of gene discovery over the past decade (*3*, *40*). This extreme etiological heterogeneity of ASD presents significant challenges to developing effective therapeutics. Identifying convergent pathways shared across genes offers a potential high-impact therapeutic target. This can be achieved through analyses of either existing datasets or those specifically generated to reflect the neurobiological context of ASD. Recent studies in this direction include work in induced neural cells, brain organoids (*41*, *42*) and assembloids (*43*, *44*), and rodent models.

Given the large numbers of genes that are implicated in ASD, large-scale screens are emerging as crucial tools to reliably detect convergent pathways. The highest throughput approaches employ CRISPR and scRNAseq, which involve substantial costs and require robust methods to jointly assess the diverse ways that gene disruption can alter the high-throughput readout. In this study, we address both the costs and the efficacy of a single RNA-seq readout to jointly assess impacts on cell fate, cell development, and intrinsic gene expression changes.

We generated interneuron progenitors relevant to understanding ASD gene mechanisms, particularly those related to potential alterations to the balance of excitation and inhibition (*53*). The expansion of the human neocortex and its increased connectivity is largely driven by an expansion of inhibitory neurons, with the increased ratio of inhibitory to excitatory neurons being specific to humans (*54–56*). Disruption of these neurons is linked to neurodevelopmental disorders and intellectual disability. In trisomy 21, an increase in neuron expression calretinin was observed (*57*), the same neuron we derived in this study. Conversely, in autism, a decrease in the density of calretinin neurons has been reported (*58*).

By leveraging an existing robust capture protocol, we demonstrate that significant cost savings are achievable with similar overall results, allowing for the use of more gRNA and deeper sequencing depth. Our results show that targeted Perturb-seq of ASD risk genes in a model of human neurodevelopment is a powerful and cost-effective approach for uncovering convergent mechanisms across the hundreds of ASD-implicated genes.

A particular challenge in neurodevelopment and other developmental processes is accounting for changes in cell fate and differentiation when interpreting gene expression changes. Previous approaches like gene expression clustering have been adapted to measure cell fate/differentiation to categorize cells into distinct clusters based on their transcriptomic profiles (*52*). This allows researchers to assess whether specific sets of sgRNAs are enriched in certain cellular subpopulations utilizing statistical methods such as chi-squared or hypergeometric distributions. In addition, specialized methods, such as differential expression (DE) analysis using tools like GSFA (*39*), MUSIC (*35*), and scMAGeCK (*36*), have been developed to address the unique characteristics of scCRISPR screens, focusing more on perturbation effects, differentially expressed genes, and the detection of gene modules.

Prior approaches are not able to simultaneously consider gene expression changes in the context of the underlying, complex cell composition dynamics. To address the limitations of previous analytical approaches, we adopted STM, integrating gene expression clustering with modeling of cell composition dynamics. By modeling gene expression changes and cell composition simultaneously, STM provides insights into the complex interplay between genetic perturbations and cellular responses in single-cell CRISPR screens. In our context, STM identifies latent topics corresponding to cell types or clusters, assigning probabilities to each cell’s topic membership. We identified 10 latent topics capturing cell type, cell fate, and stage of differentiation, making STM particularly useful for understanding the multifaceted impacts of perturbations on developmental processes.

We discovered a set of 8 ASD risk genes that, when repressed, increase the proportion of cells becoming CGE progenitors. Several genes in this set, including *ANKRD11, RFX3*, and *MBD5*, are implicated in ciliogenesis (*59–61*). As primary cilia regulate hedgehog signaling, which in turn controls ventral neuron patterning, mutations in genes affecting ciliogenesis could thereby alter interneuron fate specification (*62*). *RFX3* is expressed throughout the brain, including interneurons, and is predicted to activate the expression of other ASD genes (*63*). Zinc finger and BTB domain-containing protein 20 (*ZBTB20*) is known to bind the promoter of CoupTF1 (Nr2f1) and repress its expression (*64*), consistent with our observation of altered regionalization of ventral NPCs after *ZBTB20* repression. *KMT2A* maintains the expression *NKX2-1,* so its repression aligns with the observed shift of cells out of the *NKX2-1*+ MGE-INP cluster (*65*).

Using the STM framework, we identified four genes (*DEAF1, KMT2A, MED13L*, and *MYT1L*) with coordinated changes in topic proportions, which were replicated in an independent dataset. *MED13L*, a member of the Mediator complex, is a complex that regulates transcription and plays a role in establishing neuronal identities (*66*, *67*). Topic modulated by these genes include those related to neuronal fate commitment, consistent with the function of the Mediation complex. Recent protein-protein interaction studies have shown that the Mediator complex is a site of convergence for ASD genes (*68*). Interestingly, this study also showed that *MYT1L* interacted with the Mediator complex, and that ASD-associated mutations in *MYT1L* decreased the association between *MYT1L* and the Mediator complex. Hence, the functional relationship of MYT1L and the Mediator complex has been observed in STM analyses of our work and Li et al., in spite of different protocols and cell types, as well as on the level of protein-protein interactions, including with ASD-associated mutations. Having three studies showing related findings on the convergence of these ASD genes underscores the robust nature of this finding.

In this study, we address both the costs and the efficacy of a single RNA-seq readout to jointly assess impacts on cell fate, cell development, and intrinsic gene expression changes, and identify replicable impact of specific ASD genes on topics that reflect changes in both cell fate and neuronal differentiation. We also demonstrate significant cost savings without impacting conclusions, allowing for large scale analyses. There are, of course, further opportunities for improvement. First, while we observed significant on-target repression of target genes, employing more efficient repression systems and using multiple sgRNA per target gene could yield stronger effects (*45*, *46*). Second, maintaining the self-renewal capacity and cortical differentiation potential of iPSC-derived NPCs is an ongoing challenge (*47*) despite advances in stem cell differentiation protocols. The dorsoventral identity is a major source of variation among the differentiation of NPCs when using the dual-SMAD inhibition method (*48*).

Hybridization-based target capture after library preparation offers several advantages, including implementation on previously constructed libraries and gene panels that can be designed after pilot sequencing (*49*). While biotinylated oligos are costly, future methods (*30*) could alleviate this expense. Another limitation is that while target molecules are enriched for sequencing, they are not captured at a higher rate. TAP-seq (*29*), which employs gene-specific primers and nested PCR reactions, could serve as an alternative by enriching the actual molecules of interest earlier in the process. For Perturb-seq applications, combining both capture strategies and incorporating specific oligos for capture or reverse transcriptions could enhance sgRNA recovery (*49*). Additionally, performing multiple perturbations within the same cell could further improve scalability (*50*).

Further development of models like STM, to jointly assess impact of perturbations on multiple aspects of neurodevelopment will provide further insights. Integrating all these improvements in a 3D model of human neurodevelopment such as neural organoids or assembloids, would significantly advance the understanding of NDD and ASD risk gene function at scale (*44*, *51*).

## Acknowledgments and Disclosures

This work was supported by the BRAIN Foundation (to JDB) and with funds from the Seaver Autism Center and the Seaver Foundation. JDB also received support from the NIH (MH111679 and MH121600). MAL and XW were supported as Seaver Foundation Fellows and by NARSAD Young investigator grants (MAL 29896 & XW 31904).

CROP-seq-opti was a gift from Jay Shendure (Addgene plasmid # 106280).

This work was supported in part through the computational and data resources and staff expertise provided by Scientific Computing and Data at the Icahn School of Medicine at Mount Sinai and supported by the Clinical and Translational Science Awards (CTSA) grant UL1TR004419 from the National Center for Advancing Translational Sciences. We thank the Genomics Core Facility at the Icahn School of Medicine at Mount Sinai for performing scRNA-seq experiments, target capture, and library construction.

We are deeply grateful to Kathryn Roeder, Bernie Devlin, Nan Yang, and Silvia De Rubeis for their invaluable assistance and insightful comments throughout the preparation of this manuscript. Their expertise and feedback were instrumental in refining our study and enhancing the clarity and impact of our findings.

## Code and Data Availability

The code and pipeline used to analyze the data is available at https://github.com/xuranw/ASD_CRISPRi.

The raw from our analyses can be accessed under the XXX. The processed Seurat object is available https://github.com/xuranw/ASD_CRISPRi. The organoid data is downloaded from Zenodo (DOI: 10.5281/zenodo.7083558).

J. D.B. conceptualized the study design and supervised the project. X.W. analyzed the data. M.A.L. and U.T. generate the single-cell CRISPRi RNA-sequencing data. X.W., M.A.L., and J.B interpreted the results and drafted the manuscript. All the authors critically reviewed the manuscript and approved the final version. All the authors had full access to all the data in the study and accept the responsibility to submit for publication.

The authors report no biomedical financial interests or potential conflicts of interest.

## Notes

### Competing Interest Statement

The authors have declared no competing interest.

