## Supplementary Figures and Tables for "A scalable, high-throughput neural development platform identifies shared impact of ASD genes on cell fate and differentiation"

| **Satterstrom ASD (Fu DDD)** | **Satterstrom & Fu (GER, q < 0.1)** | | | **Satt. & Fu Neuronal** | **Fu Top 71** | **Fu DDD, syndromic** |
| --- | --- | --- | --- | --- | --- | --- |
| BCL11A | ADNP | KDM6B | SKI | GRIN2B | AUTS2 | CHAMP1 |
| CREBBP | ANKRD11 | KMT2C | SMARCA2 | NRXN1 | CTCF | MEF2C |
| FOXP2 | ARID1B | MBD5 | SMARCC2 | SCN2A | CUL3 | TSC1 |
| KMT2E | ASH1L | MED13L | SUV420H1 | SHANK2 | EBF3 |  |
| NACC1 | ASXL3 | MYT1L | TBL1XR1 | SHANK3 | EHMT1 |  |
| PPP2R5D | CHD2 | NSD1 | TCF20 | SLC6A1 | KMT2A | **X-linked** |
| RAI1 | CHD8 | PAX5 | TCF4 | SYNGAP1 | MED13 | DDX3X |
| SATB1 | CTNNB1 | PHF12 | TCF7L2 |  | NF1 | FMR1 |
| SIN3A | DEAF1 | PHF21A | TLK2 |  | NIPBL | MECP2 |
| TBR1 | DNMT3A | POGZ | TRIO |  | SATB2 |  |
| TRAF7 | DYRK1A | PTEN | VEZF1 |  | ZBTB20 |  |
| TRIP12 | FOXP1 | RFX3 | WAC |  |  |  |
|  | IRF2BPL | RORB | ZMYND11 |  |  |  |
|  | KDM5B | SETD5 |  |  |  |  |

**Supplementary Table 1:** 77 top ASD risk genes from both Satterstrom et al. and Fu et al.

| **Target Gene** | **# Cells** | **Target Gene** | **# Cells** | **Target Gene** | **# Cells** |
| --- | --- | --- | --- | --- | --- |
| ADNP | 64 | KDM6B | 192 | SATB1 | 95 |
| ANKRD11 | 256 | KMT2A* | 174 | SATB2* | 411 |
| ARID1B | 191 | KMT2C | 345 | SCN2A | 262 |
| ASH1L | 179 | KMT2E | 150 | SETD5 | 149 |
| ASXL3 | 182 | KMT5B | 304 | SHANK2 | 387 |
| AUTS2* | 201 | MBD5 | 220 | SHANK3 | 172 |
| BCL11A | 165 | MECP2* | 443 | SIN3A | 126 |
| CHAMP1* | 228 | MED13* | 538 | SKI* | 166 |
| CHD2 | 171 | MED13L | 377 | SLC6A1 | 264 |
| CHD8 | 179 | MEF2C* | 442 | SMARCA2* | 322 |
| CREBBP | 72 | MYT1L | 123 | SMARCC2 | 129 |
| CTCF* | 355 | NACC1 | 278 | SYNGAP1 | 237 |
| CTNNB1 | 67 | NF1* | 964 | TBL1XR1 | 350 |
| CUL3* | 217 | NIPBL* | 270 | TBR1 | 181 |
| DDX3X* | 266 | **NONTARGETING** | 922 | TCF20 | 279 |
| DEAF1 | 317 | NRXN1 | 317 | TCF4 | 240 |
| DNMT3A | 267 | NSD1 | 181 | TCF7L2 | 321 |
| DYRK1A | 490 | PAX5 | 171 | TLK2 | 155 |
| EBF3* | 385 | PHF12 | 67 | TRAF7 | 92 |
| EHMT1* | 220 | PHF21A | 259 | TRIO* | 322 |
| FMR1* | 408 | POGZ | 107 | TRIP12 | 243 |
| FOXP1 | 151 | PPP2R5D | 136 | TSC1* | 373 |
| FOXP2 | 44 | PTEN | 363 | VEZF1 | 253 |
| GRIN2B | 242 | RAI1 | 253 | WAC | 300 |
| IRF2BPL | 119 | RFX3 | 284 | ZBTB20* | 498 |
| KDM5B | 390 | RORB | 88 | ZMYND11* | 263 |

**Supplementary Table 2**: 77 targeted ASD risk genes, and number of cells for each of the targeted ASD risk genes. *: known ASD genes added to the Satterstrom et al list.

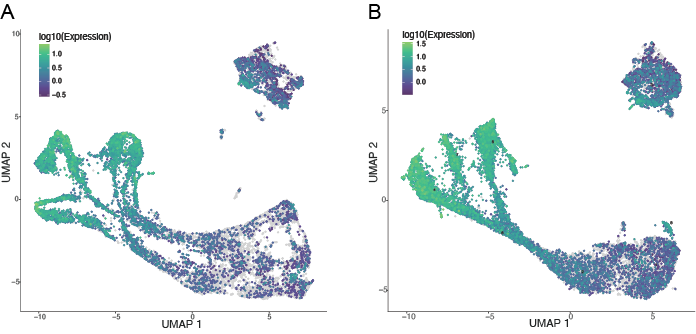

**Supplementary Figure 1:** UMAP plot from full transcriptome and targeted library. Colored by the expression level of DCX gene, the marker of neuron maturation. Two UMAP plots are generated separately. **A.** Full transcriptome with 17557 genes across 16170 cells. **B.** Targeted library with1045 genes across 17876 cells.

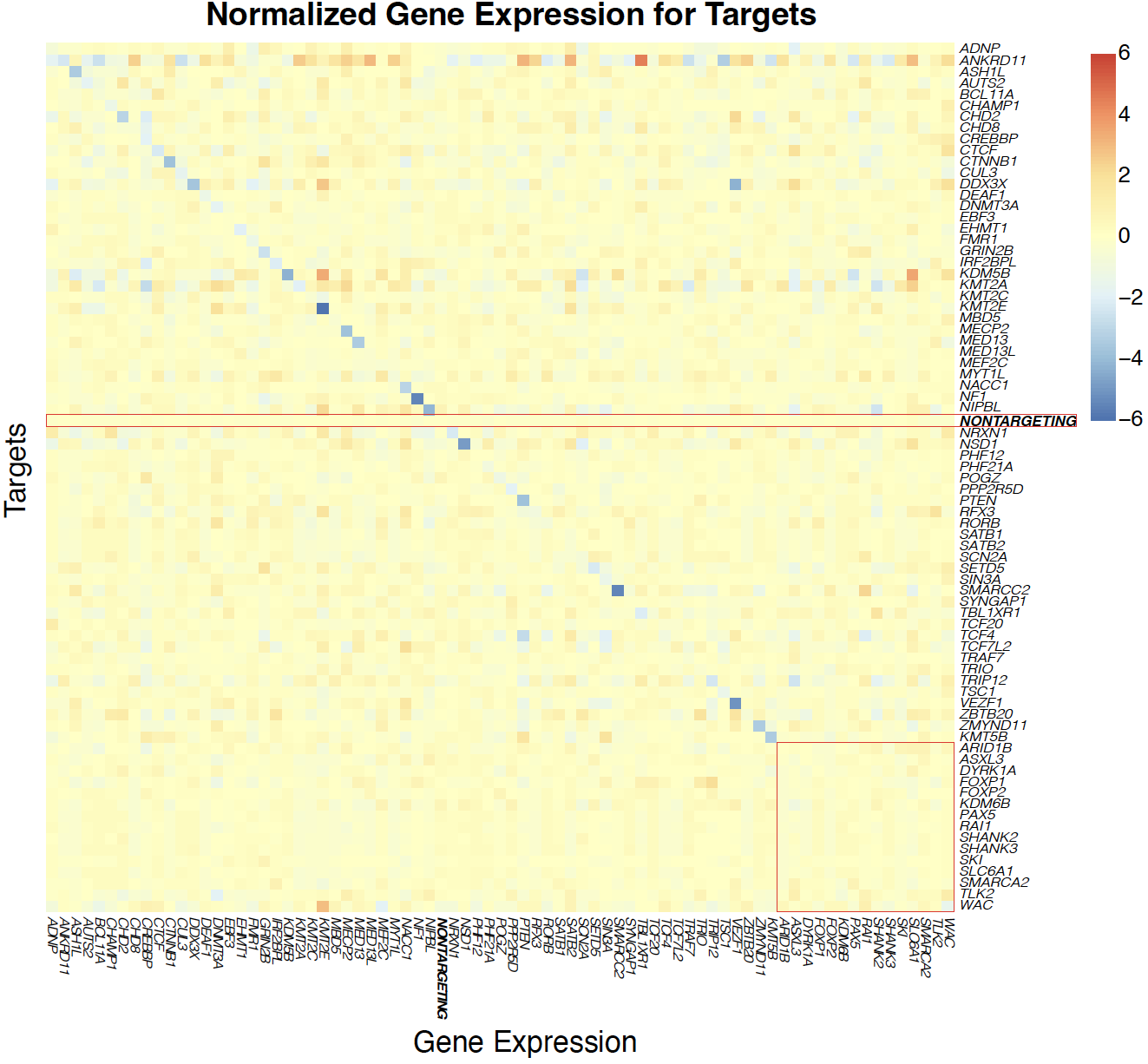

**Supplementary Figure 2.** Heatmap showing the normalized expression (Z-scores) of the targeted ASD risk genes across the CRISPRi experiments. Z-scores are calculated from the beta coefficients from MIMOSCA. Each row represents a gene expression, and each column represents a different perturbation, with the color scale indicating the gene expression level. The red square shows the targets that do not have significant effects of expression of their own. The red rectangle highlights the z-score for NONTARGETING cells.

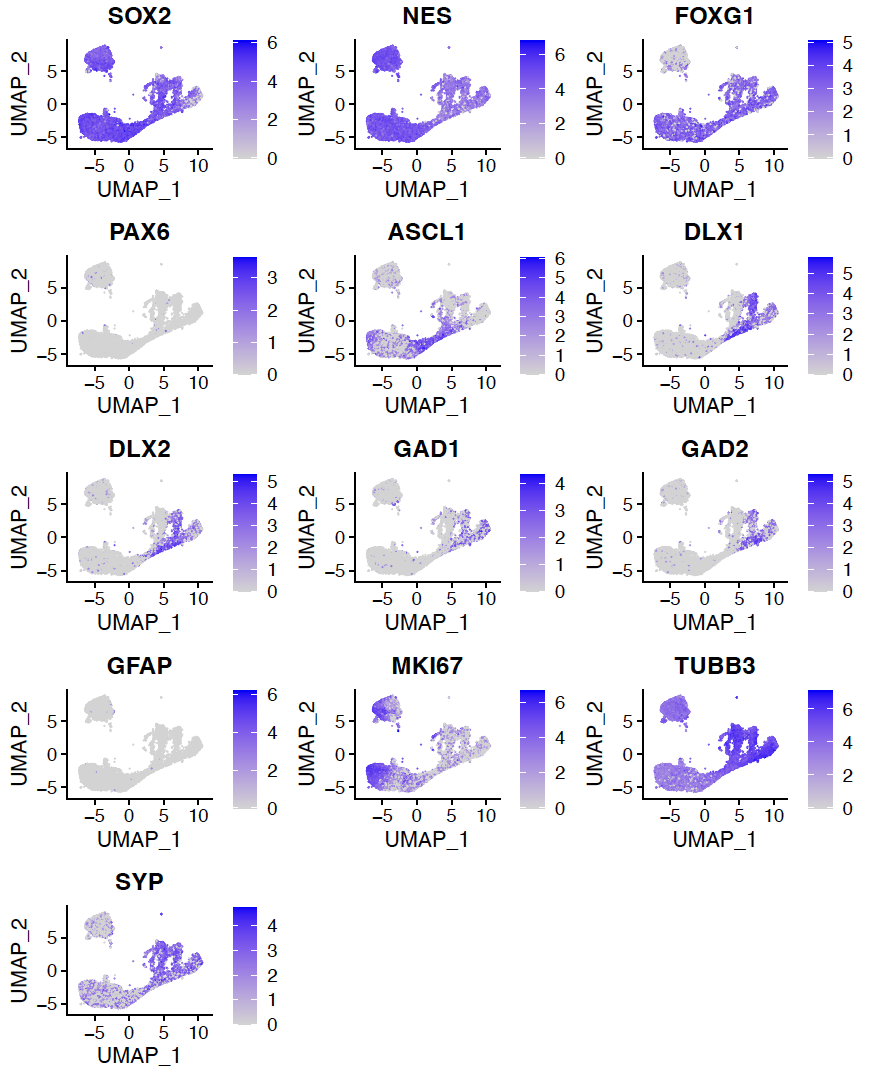
**Supplementary Figure 3**: Expression of marker genes colored on UMAP. Forebrain neural progenitors: *SOX2, NES, FOXG1*. Dorsal cortical markers: *PAX6* and *EMX2*. Ventral fate: *ASCL1* and *DLX2*. Gamma-aminobutyric acid-containing (GABAergic) neuron markers: *GAD1, GAD2*, and *vGAT*. Developmental marker gene expression: *MKI76, TUBB*, and *SYP*.

| Target | p-val | Adj-p-val | Target | p-val | Adj-p-val | Target | p-val | Adj-p-val |
| --- | --- | --- | --- | --- | --- | --- | --- | --- |
| ADNP | 0.738 | 1 | KDM5B | 0.034 | 1 | RFX3 | 0 | 0 |
| ANKRD11 | 0 | 0 | KMT2A | 0 | 0 | RORB | 0.467 | 1 |
| ASH1L | 0.302 | 1 | KMT2C | 0 | 0.004 | SATB1 | 0.429 | 1 |
| AUTS2 | 1 | 1 | KMT2E | 0.844 | 1 | SATB2 | 0.001 | 0.043 |
| BCL11A | 0.144 | 1 | MBD5 | 0 | 0 | SCN2A | 0.369 | 1 |
| CHAMP1 | 0.612 | 1 | MECP2 | 0.937 | 1 | SETD5 | 0.703 | 1 |
| CHD2 | 0.977 | 1 | MED13 | 0.016 | 0.963 | SIN3A | 0.873 | 1 |
| CHD8 | 0 | 0.008 | MED13L | 0 | 0 | SMARCC2 | 0.308 | 1 |
| CREBBP | 0.021 | 1 | MEF2C | 1 | 1 | SYNGAP1 | 0.132 | 1 |
| CTCF | 0.699 | 1 | MYT1L | 0 | 0 | TBL1XR1 | 0.007 | 0.449 |
| CTNNB1 | 0.001 | 0.05 | NACC1 | 0.018 | 1 | TCF20 | 0.094 | 1 |
| CUL3 | 0.038 | 1 | NF1 | 0.114 | 1 | TCF4 | 0 | 0 |
| DDX3X | 0.1 | 1 | NIPBL | 0 | 0.007 | TCF7L2 | 0 | 0 |
| DEAF1 | 0 | 0 | NRXN1 | 0.482 | 1 | TRAF7 | 0.029 | 1 |
| DNMT3A | 0.01 | 0.593 | NSD1 | 0.715 | 1 | TRIO | 0.401 | 1 |
| EBF3 | 0.003 | 0.206 | PHF12 | 0.615 | 1 | TRIP12 | 0.003 | 0.181 |
| EHMT1 | 1 | 1 | PHF21A | 0.235 | 1 | TSC1 | 0.056 | 1 |
| FMR1 | 0.32 | 1 | POGZ | 0.006 | 0.338 | VEZF1 | 0.746 | 1 |
| GRIN2B | 0.512 | 1 | PPP2R5D | 1 | 1 | ZBTB20 | 0 | 0 |
| IRF2BPL | 0.087 | 1 | PTEN | 0.033 | 1 | ZMYND11 | 0.556 | 1 |

**Supplementary Table 3**: The raw p-value from chi-square test and the adjusted p-value. Significant targets are highlighted and the cut-off = 0.01.

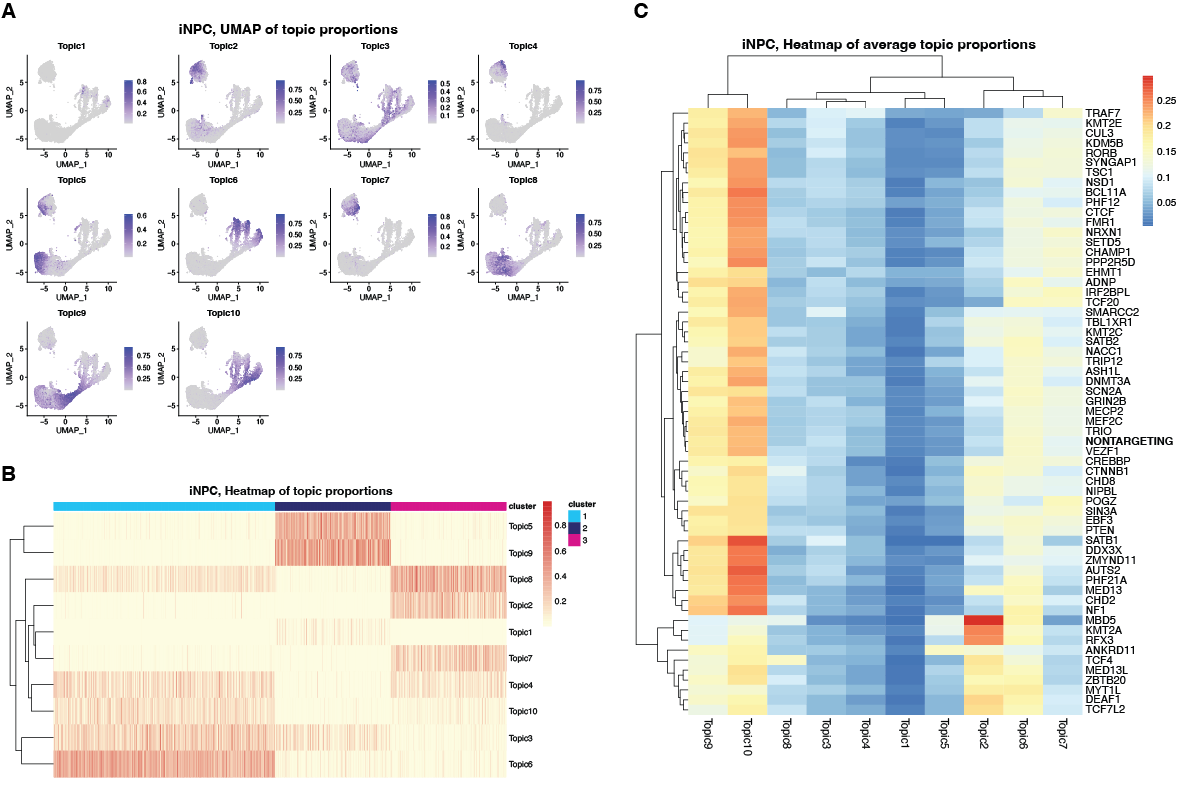

**Supplementary Figure 4**: Estimated topic proportions from iNPC data using structural topic model (STM). **A.** The UMAP of estimated topic proportions for each topic; **B.** The heatmap of estimated topic proportions, colored by estimated topic proportions. Cells are ordered by the broad cluster defined by marker genes and UMAP plots; **C.** The heatmap of the average topic proportions

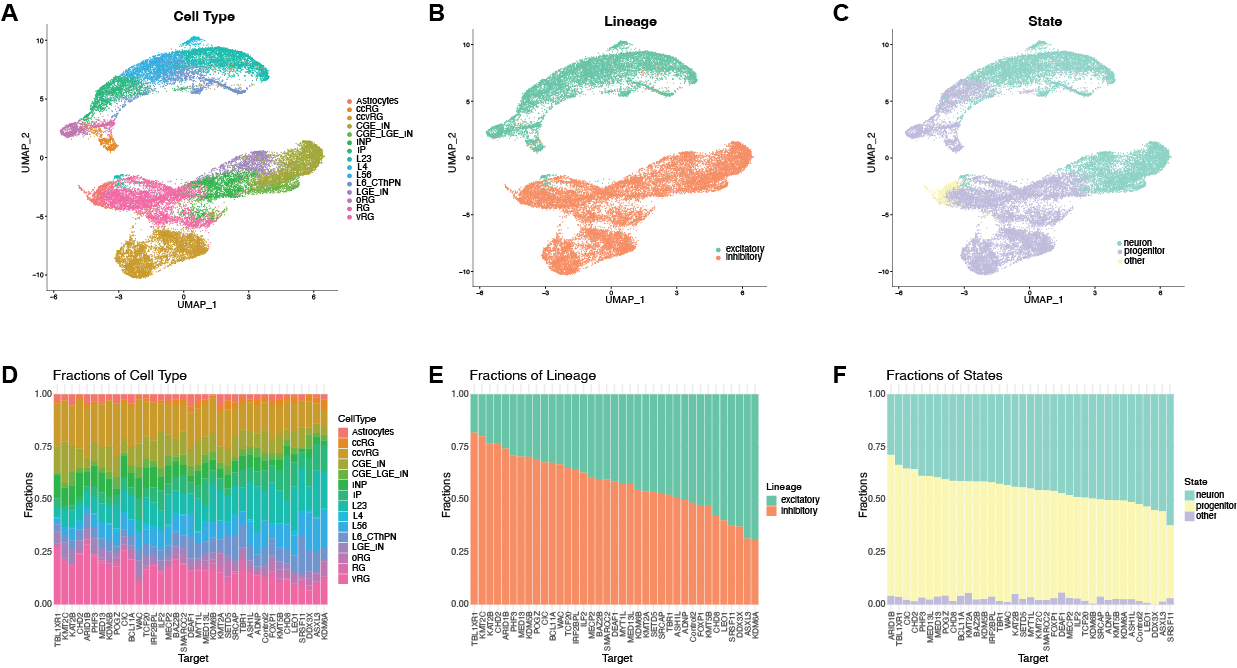

**Supplementary Figure 5**: Summary of the organoid dataset. **A-C.** UMAP of organoid dataset colored by **A.** pre-annotated cell types; **B.** Lineage of excitatory neurons and inhibitory neurons; **C.** State of progenitor, neurons and other cells (glia cells). **D-F.** Bar plot of targets and control cells from the organoid dataset colored and ordered by the fractions of **A.** pre-annotated cell types; **B.** Lineage of excitatory neurons and inhibitory neurons; **C.** State of progenitor, neurons, and other cells (glia cells).

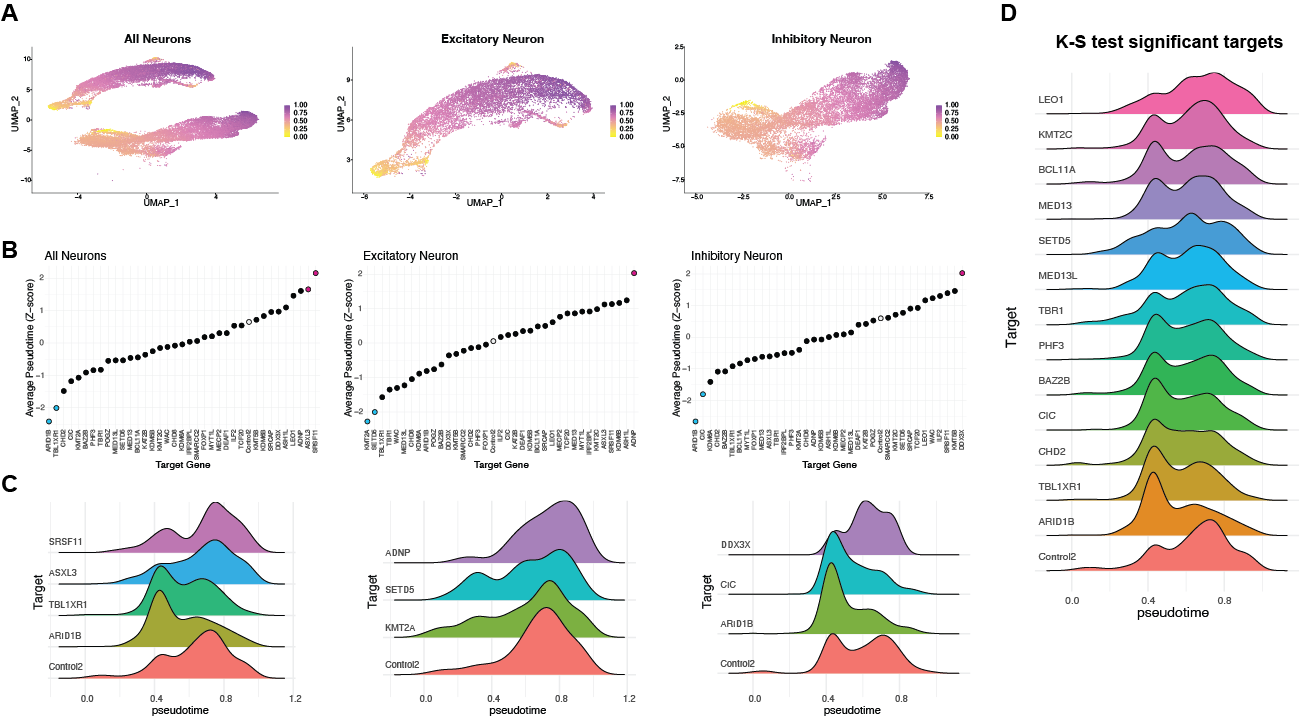

**Supplementary Figure 6**: Pseudotime analysis of all neuron cells (remove the glia cells) from the organoid dataset. **A.** UMAP of cells colored by pseudotime. The three panels correspond to all neuron cells, excitatory neurons and inhibitory neurons. **B.** Z-score pseudotime grouped by targets for all neuron cells and two lineages. Significant changes in z-scores are determined by Norm(0.95) and colored yellow and purple respectively. **C.** Density plot of the pseudotime for control cells and the targeted cells that show significant pseudotime density shift in all neurons and in two lineages. Significant targets are determined by the z-scores in B. **D.** Density plot of the pseudotime for control cells and significant targeted cells for all neurons. The significant targets are determined by Kolmogorov-Smirnov test with p-value < 0.05.
